# Iteratively improving Hi-C experiments one step at a time

**DOI:** 10.1101/287201

**Authors:** Rosela Golloshi, Jacob Sanders, Rachel Patton McCord

## Abstract

The 3D organization of eukaryotic chromosomes affects key processes such as gene expression, DNA replication, cell division, and response to DNA damage. The genome-wide chromosome conformation capture (Hi-C) approach can characterize the landscape of 3D genome organization by measuring interaction frequencies between all genomic regions. Hi-C protocol improvements and rapid advances in DNA sequencing power have made Hi-C useful to diverse biological systems, not only to elucidate the role of 3D genome structure in proper cellular function, but also to characterize genomic rearrangements, assemble new genomes, and consider chromatin interactions as potential biomarkers for diseases. Yet, the Hi-C protocol is still complex and subject to variations at numerous steps that can affect the resulting data. Thus, there is still a need for better understanding and control of factors that contribute to Hi-C experiment success and data quality. Here, we evaluate recently proposed Hi-C protocol modifications as well as often overlooked variables in sample preparation and examine their effects on Hi-C data quality. We examine artifacts that can occur during Hi-C library preparation, including microhomology-based artificial template copying and chimera formation that can add noise to the downstream data. Exploring the mechanisms underlying Hi-C artifacts pinpoints steps that should be further optimized in the future. To improve the utility of Hi-C in characterizing the 3D genome of specialized populations of cells or small samples of primary tissue, we identify steps prone to DNA loss which should be optimized to adapt Hi-C to lower cell numbers.

**Highlights:** 3 to 5 bullet points (maximum 85 characters, including spaces, per bullet point)

- **Variability in Hi-C libraries can arise from early steps of cell preparation**
- **Hi-C 2.0 changes to interaction capture steps also benefit 6-cutter libraries**
- **Artificial molecule fusions can arise during end repair and PCR, increasing noise**
- **Common causes of Hi-C DNA loss identified for future optimization**

## 1. Introduction

The billions of DNA bases in the eukaryotic genome must be efficiently packed to fit inside the micron-sized nucleus and to perform necessary cellular functions including gene expression as well as DNA replication and repair [1-3]. Abnormal 3D packaging of chromatin may disrupt cellular homeostasis and lead to diseases [4, 5].

Many techniques have been devised to understand how chromatin is packed into the nucleus and how that packaging affects genome function. Light and electron microscopy led to observations of densely packed heterochromatin and open euchromatin as well as the small scale wrapping of DNA around nucleosomes [6, 7]. Fluorescence microscopy approaches such as fluorescence *in situ* hybridization (FISH) [8] and *in vivo* techniques such as bacterial operator arrays [9] and CRISPR imaging [10] have allowed visualization and tracking of the nuclear arrangements of specific genome loci. In recent years, the development of the chromosome conformation capture (3C) technique [11], and variations such as 4C [12, 13], 5C [14], and Hi-C [15], have accelerated the progress of the genome organization field and sparked the interest of researchers with numerous biological problems and across disciplines from genetics to physics, mathematics, and computer science.

As high throughput sequencing becomes increasingly powerful and inexpensive, many researchers have adopted Hi-C as a method to capture a snapshot of 3D nuclear organization genome-wide. The ability to represent 3D genome organization in terms of matrices of contact frequencies allows the characterization of structures across a variety of length scales, from the positioning of whole chromosome territories to the folding of small-scale enhancer-promoter loops [3, 16, 17]. Even when more focused resolution is desired, many techniques begin with an approach similar to Hi-C and then isolate a subpopulation of interactions of interest. This is the case with approaches such as Capture-C [18], Hi-ChIP [19], and others. Further, it has recently become clear that Hi-C is a very useful tool not only for measuring 3D genome folding, but also for *de novo* whole genome sequence assembly [20-22] and translocation detection [23]. As basic researchers from many fields, commercial ventures (such as Phase Genomics and Dovetail Genomics), and clinical researchers adopt Hi-C based techniques, it is more important than ever to optimize the methodological details of the Hi-C protocol and understand how they affect the resulting data and interpretation.

Several excellent discussions of Hi-C theory and optimizations of Hi-C protocols have been published in recent years [17, 24-26], and numerous other insights about key steps of the Hi-C protocol have been discovered while researchers optimize this technique in various systems. Yet, Hi-C is still a technique that often requires troubleshooting and can sometimes unexpectedly result in low quality datasets even when the best current protocols are followed. Rather than a comprehensive guide or update to every step of the Hi-C approach, here, we will describe our observations about certain key variables in the Hi-C protocol and the resulting data. We will present evidence about the effect of certain protocol modifications on Hi-C data quality and utility, propose explanations for previously unreported artifacts that can arise in Hi-C libraries, describe considerations for applying Hi-C to smaller populations of cells, and discuss key areas of uncertainty that will be ripe candidates for future careful optimization.

## 2. Early steps in the Hi-C protocol with downstream implications

### 2.1 Initial sample preparation: an overlooked variable

The key initial steps of Hi-C involve the covalent crosslinking of chromatin regions interacting in 3D space, followed by restriction enzyme digestion, marking digested ends with biotin, and proximity ligation of fragments from interacting regions. As we will discuss below, digestion, biotin fill-in, and ligation conditions certainly can have a large impact on Hi-C library quality, but another major source of variation in Hi-C is the handling of the starting material before restriction enzyme digestion takes place. For cells in culture, variables such as exactly how the formaldehyde is added, how adherent cells are removed from culture dish, and how nuclei are isolated and permeabilized can have downstream implications for Hi-C library quality (Figure 1A).

**Figure 1.**
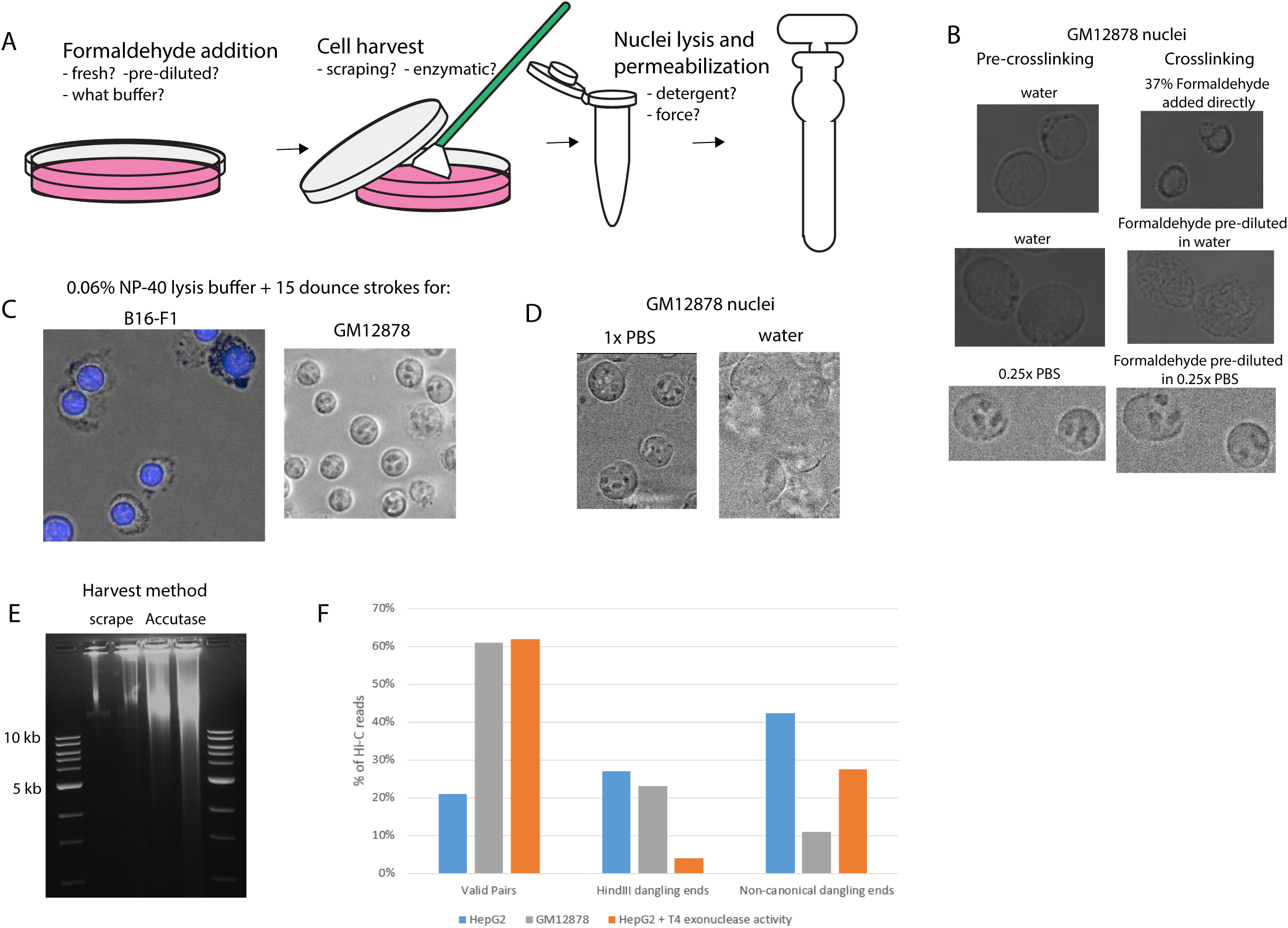
Variable factors in early steps of the Hi-C protocol. A) Hi-C experiments for cells in culture begin with decisions about initial steps such as formaldehyde crosslinking, cell harvest, and nuclei lysis and permeabilization. B) Different approaches to adding formaldehyde may affect the state of the nucleus. In each case, the final concentration of formaldehyde is 1%. (Top) Nuclei directly under the site of 37% formaldehyde addition show a dramatic shape change. (Middle) Nucleus size change is less apparent when formaldehyde is pre-diluted to 1% in water, but nuclear wrinkling is still apparent. (Bottom) Nuclear structure is preserved when formaldehyde is pre-diluted in 0.25x PBS. C) B16-F1 mouse melanoma cells retain extensive cytoplasmic debris in the same nucleus purification protocol that effectively purifies GM12878 nuclei. D) GM12878 nuclei can lyse when 1x PBS buffer is replaced with water, indicating that varying buffer conditions could contribute to higher or lower rates of intact nuclei entering the downstream Hi-C protocol. E) The appearance and quantity of DNA varies for HepG2 cells harvested by either scraping after crosslinking or enzymatic removal followed by crosslinking. In each case, crosslinks were reversed in harvested cells for 1 hour at 65°C with proteinase K and then DNA was run on a 0.8% agarose gel. F) Hi-C protocols performed simultaneously on two different cell types can result in differing proportions of informative interaction pairs (Valid Pairs). While unligated HindIII fragments (HindIII dangling ends) are consistent between the cell types, one experiment suffered from many more random breaks or biotin incorporation at nicks that result in non-canonical dangling ends. These non-canonical ends are not reduced as much as HindIII dangling ends are reduced by additional biotin removal from DNA ends with the exonuclease activity of T4 DNA polymerase.

The development of the “in situ”[17] or “in nucleus”[27] Hi-C protocol, where nuclei are isolated and permeabilized, but not lysed, before digestion and ligation, acknowledges the importance of the initial state of the biological material at the beginning of Hi-C. Avoiding high concentrations of SDS, nuclei lysis, and dilution to large volumes during digestion and ligation increases Hi-C reproducibility and decreases the capture of random background interactions [17, 27]. The generally accepted explanation for this improvement has been that intact nuclei constrain the movement and random collisions of crosslinked complexes, but other factors could be at work as well. Other work has suggested that high concentrations of SDS and subsequent Triton sequestration of SDS are more likely to lead to aggregates of material that reduce digestion, fill-in, and ligation efficiency [28].

We have evidence that suggests that not only nuclear lysis, but also the random chromatin damage that may accompany physical forces that lyse nuclei, may also contribute to variations in Hi-C protocol success. Unfortunately, even when a theoretically consistent protocol is used, there can be variation in what actually happens to a given cell type during the initial steps of Hi-C.

#### 2.1.1 Formaldehyde Crosslinking

The effects of variations in formaldehyde concentration and crosslinking time on Hi-C results have been considered previously [29, 30]. It has also been noted that the presence of serum in the cell culture media can inhibit effective crosslinking [31]. Other work has suggested that the preparation of fresh formaldehyde immediately before each experiment, to prevent uncontrolled formaldehyde polymerization, is helpful to increase reproducibility [28]. In addition to these previous observations, we have observed that dramatic changes in the appearance of nuclei can occur depending on how formaldehyde is added (Figure 1B). When concentrated formaldehyde (37%) is added directly to isolated nuclei, an instantaneous and irreversible shrinkage of nuclei nearest to the site of formaldehyde addition is observed. Notable, though less dramatic, alterations in nucleus appearance were also observed when formaldehyde pre-diluted to 1% in water was added to nuclei already incubated in water. The best preservation of pre-existing nuclear appearance was achieved by replacing cell culture media with a pre-mixed 1% solution of formaldehyde in either serum-free cell culture media, or other buffered solutions (PBS, HBSS, HEPES). Thus, we recommend that formaldehyde be pre-diluted in an appropriate buffer before addition to the cells destined for Hi-C.

#### 2.1.2 Cell Harvest

The harvest of crosslinked cells is a common step to many genomic experiments (including ChIP-Seq and others), but there are still considerations in this step that are often unexplored. We have observed that different cells types vary greatly in how easily they can be recovered by manual scraping from culture dishes after crosslinking. Differentiated myotubes can require trypsinization to aid release from the dish [32] while colony forming cells like HepG2 detach very easily (unpublished observations). Consequently, different cellular preparations may unintentionally be treated more or less harshly during cell scraping, even by different individuals performing the same protocol. One option for increasing the reproducibility and recovery of material is to gently enzymatically detach cells from the plate using, for example, Accutase, before crosslinking and then to perform crosslinking steps in a single cell suspension. Our results indicate that this approach can lead to better estimation of cell numbers going into the experiment, less cellular aggregates that may be hard to permeabilize, and higher recovery of DNA from crosslinked cells (Figure 1E). But, since cellular attachments to different substrates can affect nuclear structure [33], cells may not retain their native 3D genome conformation after being detached. So, crosslinking attached cells in their native state is often preferable, and scraping conditions must be carefully considered and documented for each new cell type and condition. Crosslinked plates should be evaluated under the microscope to determine the minimal amount of scraping needed to retrieve the majority of the cells.

#### 2.1.3 Nuclei Isolation / Permeabilization

Most Hi-C protocols recommend that nuclei be released by dounce homogenization before restriction digestion begins. But, like cell scraping, this technique is inherently variable depending on how fast or slow a given individual performs the recommended number of strokes, or the variable properties of individual cells. Indeed, we have observed that different cell types can look quite different after the same nuclei isolation steps. While GM12878 nuclei are cleanly isolated after incubation in a detergent-containing buffer and 15 strokes of the dounce homogenizer, mouse melanoma cells still retain a large amount of debris around their nuclei after the same treatment (Figure 1C). Relatedly, changes in buffer conditions can cause dramatic effects on isolated nuclei, again, depending on cell type. We have observed that upon addition of 1 drop of pure water to isolated nuclei from GM12878 cells, the nuclei can dramatically lyse (Figure 1D), while a HeLa epithelial-type cancer cell nucleus is minimally affected by the same treatment. In practice, variations are often noted in the visible properties of Hi-C nuclei before the digestion step: some nuclei settle quickly or are more clumpy, granular, or sticky. But such differences are rarely reported or considered. To develop more careful understanding of what upstream steps lead to these differences, and how they affect downstream steps, during the optimization of Hi-C for a new cell type or by a new experimentalist, we recommend regular microscopic visualization of cells and nuclei for better cataloging of their state after different manipulations.

#### 2.1.4 Evidence that variable DNA damage affects Hi-C library quality

Poorly controlled variations in the steps described above can have a negative impact on the downstream quality of the Hi-C library. Insufficient cell recovery during cell harvest steps can lead to low amounts of DNA and low library complexity. Incomplete nucleus isolation or permeabilization can affect the efficiency of downstream steps in which enzymes must access the chromatin. As previously described, DNA recovery and digestion efficiency can and should be checked on a gel before subsequent steps [24, 31]. But, it is more difficult to assay early in the protocol whether the cells have been treated too harshly, leading to complete nuclear lysis or chromatin damage. In both long read Sanger sequencing of isolated Hi-C library molecules and high throughput sequencing results, we have observed that different replicates and cell types can have varying proportions of breaks in the DNA at sites that do not correspond to restriction enzyme sequences (discussed in detail as “random breaks” in Imakaev et al. [34]). Some of these non-canonical breaks correspond to unligated but biotinylated molecules that end up being uninformative reads (“dangling ends”) at the end of the Hi-C experiment, reducing the number of interaction pairs recovered. These non-canonical dangling ends also seem to be recalcitrant to attempts to remove biotin at fragment ends (Figure 1F), and may therefore arise from biotin incorporation at nicks in the middle of fragments. A higher proportion of random breakage also often corresponds to higher noise levels (manifested as a higher background of interchromosomal interactions). Overall, it is important to consider and take steps to avoid the factors above that may increase unintentional nucleus and chromatin damage.

#### 2.1.5 Future optimization of cell preparation needed

All of the variables discussed apply in particular to Hi-C being performed on cells in culture. Controlling the consistent initial preparation of crosslinked material before digestion is even more challenging when whole primary tissues are the source of biological material. Numerous upstream approaches to prepare diverse biological materials have been developed [20, 35-37], but many of these require similar downstream steps discussed above. To increase the success of the increasingly complex preparations of diverse biological materials for future Hi-C experiments, future work should explore protocols that reduce variability in ambiguous steps such as scraping and douncing. There is also a great need for more informative quality control tests of initial Hi-C steps that will allow researchers to assay whether a given preparation of material is suitable (good chromatin accessibility and material recovery with low DNA damage and nuclei lysis) for the expensive downstream steps of digestion, biotin fill-in, and ligation, not to mention library preparation and sequencing.

### 2.2 Utility of Hi-C 2.0 conditions for HindIII libraries

Recently, Belaghzal et al. have developed an optimized protocol called Hi-C 2.0 which uses the DpnII enzyme and is designed for single aliquots of 5 million cells [24]. Some changes in the protocol from previously published approaches [17, 31] might seem to be minor or just specific to the use of biotin-dATP and DpnII. However, we find that these variations also notably improve HindIII Hi-C library quality in a direct comparison using aliquots of 5 million mouse melanoma cells (B16-F1) identically prepared in initial steps (see Appendix A for protocols used). In particular, the modified protocol involves performing biotin-dCTP fill-in at 23°C for 4 hours instead of 37°C for 2 hours. When applied to a HindIII Hi-C library preparation, these modifications resulted in a nearly 10% increase in valid interaction pairs due to a dramatic decrease in unligated dangling ends (Figure 2A). Notably, the decrease in dangling end molecules corresponded to a reduction of the bias toward inward facing reads among valid pairs. This indicates that a fraction of the excess inward facing interaction pairs are actually undigested dangling ends, which are then also reduced when dangling ends are removed.

**Figure 2.**
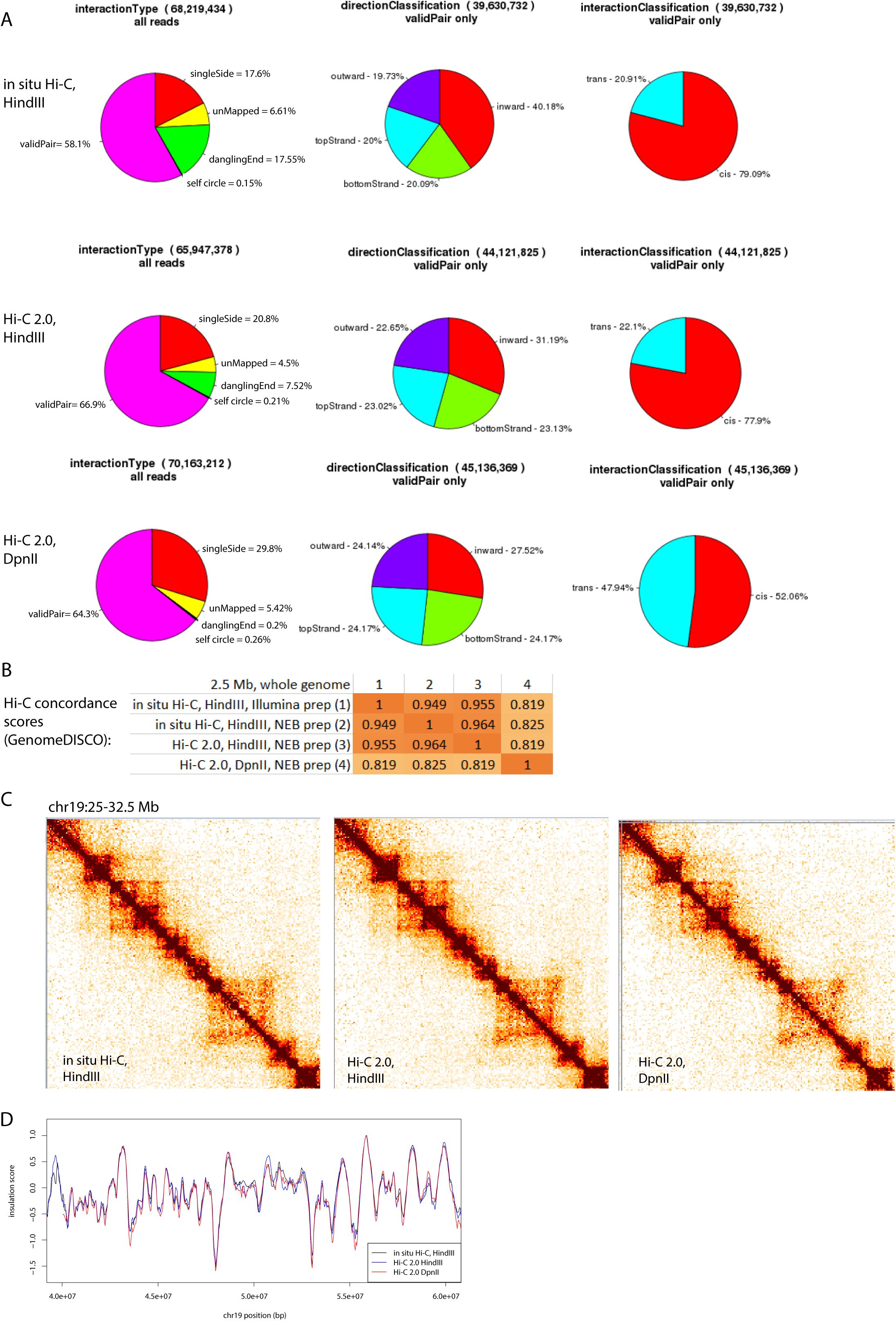
Hi-C 2.0 [24] protocol changes benefit HindIII Hi-C libraries as well as DpnII libraries. A) Hi-C library statistics for aliquots of 5 million B16-F1 mouse melanoma cells prepared with: in situ Hi-C with HindIII (top), Hi-C 2.0 with HindIII (middle), and Hi-C 2.0 with DpnII (bottom). Left: proportions of reads that are valid interaction pairs as compared to unmapped or single fragment reads. Middle: proportions of mapping orientations of valid interaction pairs. Hi-C 2.0 gives less inward bias, indicating a reduction of biases arising from undigested fragments. Right: Fraction of intrachromosomal (cis) and interchromosomal (trans) interaction pairs. B) Concordance between interaction maps resulting from different Hi-C protocol approaches as calculated by GenomeDISCO. C) Similarity of 40 kb binned and iteratively corrected [34] heatmaps from the 3 parallel experiments for a 7.5 Mb region of chr19. D) Insulation score profiles (sliding window size of 500 kb, heatmap resolution of 40 kb) for a section of chromosome 19 for each Hi-C protocol. Overall TAD boundaries (major dips in the insulation score) are conserved in position and strength between the Hi-C protocols at this resolution.

Dangling ends are reduced even further when DpnII with biotin-dATP is used instead of HindIII in the Hi-C 2.0 protocol (Figure 2A). However, an increase in interchromosomal (“% trans”) interactions is observed, which could indicate higher background noise in this condition. This increase in the fraction of interchromosomal interactions when the cells were derived from the same initial preparation steps shows that nuclear lysis (which would be the same in each of these aliquots) is not the only factor that can contribute to higher levels of potential background ligation. Overall, all of these approaches lead to reproducible final Hi-C libraries according to concordance scores calculated by GenomeDISCO [38] and visual comparisons of the resulting heatmaps at 40 kb resolution (Figure 2B and C). TAD boundary locations and strengths at 40 kb resolution are also overall similar according to an insulation score profile (Figure 2D) [37]. Thus, the major gain in these particular optimizations will lie primarily in the fraction of useful reads gained from a certain number of cells and improvements in heatmaps at higher resolution.

## 3. Approaches and Artifacts during Hi-C library preparation for sequencing

After the “3C-like” phase of the Hi-C experiment is complete (digestion, biotin fill-in, and ligation), the Hi-C DNA follows some steps that are common to nearly all high throughput sequencing experiments but with variations and factors unique to Hi-C that require special consideration. Even with a high quality library at the beginning of this stage, variations in library preparation can matter, and problems with low quality libraries become exacerbated during preparations for sequencing.

### 3.1 Reproducible NEBNext library preparation of captured interactions

Like some previously published protocols [17], we find it useful to perform streptavidin bead capture of biotinylated interactions immediately after size selection and before any end repair. We have adapted the NEBNext® Ultra™ II DNA Library Prep protocol for use on beads, resulting in lower reaction volumes, fewer steps, and reduced time than some Hi-C protocols (Appendix A). In our hands, this protocol is easier and more reproducible specifically in cases of indexed adaptor ligation than adapting the Illumina protocol to Hi-C, possibly due to the divergent design of NEB adaptors. We find that substantial dilution of adaptors (at least 1:10) is necessary to avoid excess adaptors in the final Hi-C library.

### 3.2 Disadvantages of over-sonication to short DNA fragment length

Given that ligated interacting DNA molecules are often multiple kilobases in length, fragmentation of DNA to smaller sizes for sequencing on the Illumina platform is essential. However, some evidence suggests that it is better to err on the side of leaving larger DNA fragments rather than trying to achieve average sizes of 200 bp, as has sometimes been recommended [24, 31]. Smaller fragments reduce the overall mappability of the Hi-C library after sequencing, in part because shorter sequences are less likely to be uniquely mappable, particularly in repetitive regions. In Hi-C specifically, short fragments are also likely to contain more reads if one of the two sides of the interacting pair is too short to be mapped before the sequencing runs into the junction with the other interacting partner (Figure 3A). This phenomenon results in a higher proportion of “single side mapped” reads. Also, evidence further discussed below suggests that shorter molecules are more likely to contain end-repair related artifacts (Figure 3F).

**Figure 3.**
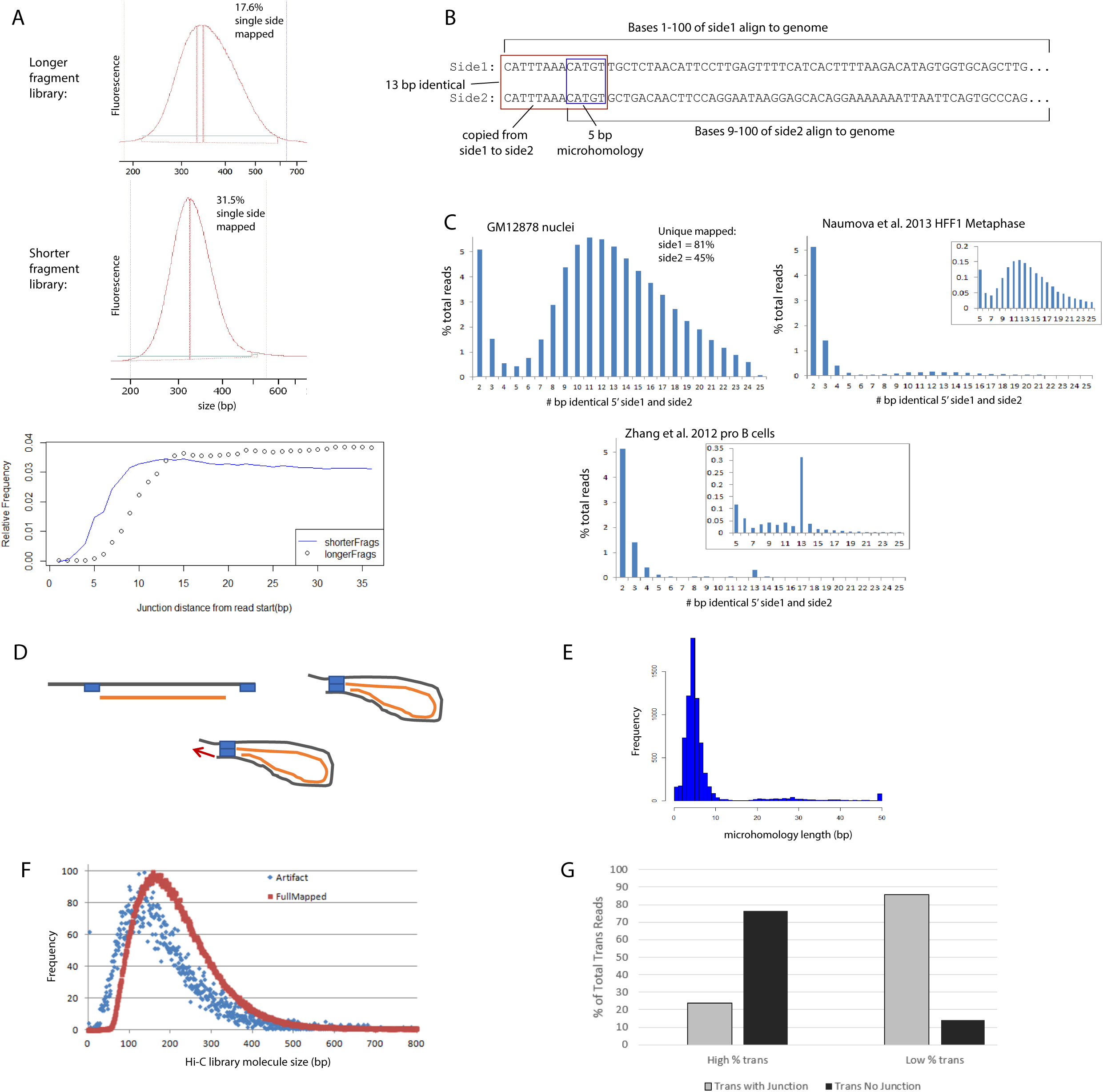
Concerns and artifacts arising from Hi-C sequencing library preparation. A) Fragment size distributions of the final Hi-C library (ligated DNA insert with ligated adaptors) from an Agilent Bioanalyzer trace. The shorter fragments with a narrower distribution have a higher percentage of single side mapped reads. This is likely attributable to the higher chance of encountering a ligation junction in the first few base pairs of the shorter fragment (blue line, bottom panel), resulting in one unmappable interaction partner. B) Sample reads demonstrating the artifact in which a piece of side1 is artificially attached to side2. As a result, side1 and 2 are identical for the first 10-15 bp and side2 is unmappable without 5’ trimming. C) A Hi-C experiment on GM12878 nuclei showed an extreme case of side1 copying onto side2, revealed as a large % of the total read pairs with 10-20 bases identical between side1 and side2, resulting in only 45% alignment rate of side2 reads. This effect is observed to a lesser, but notable extent in other experimental data, such as published HFF1 metaphase Hi-C [29]. Other published datasets, such as pro B cell Hi-C data [3] show only a peak at 12 bp, which corresponds to the amount of identity between adapter dimers. D) Model of explanation of how short homologous regions (“microhomologies”, blue boxes) could form hairpins and then elongate during end repair, copying the rest of side1 (left end) onto side2 (right end). E) Most read pairs experiencing this copy artifact have 4-6 bp of identical sequence at their mapped ends that could form hairpins. F) Reads containing these artifacts (blue) tend to arise from shorter DNA molecules than non-artifact reads (red). G) In a Hi-C library with large numbers of noisy interchromosomal read pairs (“High % trans”), a high percentage of these reads contained no Hi-C junction sequence, suggesting that they may arise from something other than ordinary Hi-C ligation. When the number of interchromosomal pairs is low, on the other hand, most of these reads contain a canonical Hi-C junction sequence.

### 3.3 Dangers of chimera formation and end repair based artifacts

In all high throughput genome-wide sequencing approaches, the complexity of DNA sequences in the same tube raises the possibility that small regions of homology between molecules could hybridize and form chimeras during PCR amplification. However, in techniques other than Hi-C, chimeric reads can easily be excluded due to their impossible genomic positions (paired reads from different chromosomes, reads from the same DNA strand, etc.). In Hi-C, by contrast, informative paired sequence reads should come from all orientations and locations in the genome, making these possible artifacts harder to detect or exclude.

We have found evidence of one particular type of artifact that shows the possibility of such homology-based chimeras. In the sequencing results of some Hi-C experiments, we found an unusually low rate of mapping on side2 of the paired end sequencing results (81% mappability for side1 vs. 45% for side2). Strangely, we discovered 10-15 bases of side1 had been copied artificially onto the beginning of the side2 read, and thus side2 would not map unless the first bases were excluded (Figure 3B and C). Further investigation revealed that molecules with this artifact had 4-6 bp of microhomology between the mappable ends of side1 and side2 (Figure 3E). This artifact is most common in libraries that have undergone extensive biotin removal steps and sonication to small fragment sizes (Figure 3F), both conditions that would tend to expose single stranded overhangs. During the room temperature end repair step, hairpins could form between the short complementary single stranded ends, and the T4 DNA polymerase would be able to elongate the DNA from this hairpin “priming”, copying the end of side1 onto side2 (Figure 3D). Microhomologies leading to pairing between the ends of side1 and side2 fragments may be further reinforced by homology between adapter sequences on side1 and side2. Indeed, in many sequencing libraries, the only evidence for exact matches between side1 and side2 sequence reads comes from adapter dimers (Figure 3C).

Hairpins resulting from a few identical bases may sound unbelievable, but, interestingly, a very similar mechanism is known to happen *in vivo* during certain types of DNA repair [39]. If there are 3’ single stranded overhangs on damaged DNA ends, microhomologies of 1-4 bp can anneal and undergo fill-in synthesis by DNA polymerase [40, 41]. Microhomology annealing can be a small bias in non-homologous end joining repair or microhomology mediated repair pathways. As seen in Hi-C, this mode of DNA repair can lead to genomic rearrangements.

This artifact is minor in most Hi-C libraries, but suggests that care must be taken not to over-sonicate the DNA or to overdo the biotin removal recessing of DNA ends. Future library preparation protocols could also consider variations to the end repair steps to avoid the possibility of low temperature priming and extension from small identical regions. The use of tagmentation for adapter incorporation, as has already been used for some 3C protocol variants [19, 36], would likely reduce the risk of this kind of artifact.

Fortunately, the direct copying of side1 onto side2 is easily detectable, but more difficult to detect chimeras could form in a similar way either during the end repair or final PCR steps. As previously documented [31], concatenations of Hi-C library molecules cause an upward shift in overall Hi-C library molecular weight after too many rounds of PCR amplification. But, such chimeras may form before this shift is visibly detectable. Indeed, in libraries with more noisy random interchromosomal (“trans”) ligations, a higher proportion of these interchromosomal interaction reads lack the canonical Hi-C junction sequence (Figure 3G). Such random interaction noise without true ligation junctions could arise from PCR chimeras formed in an attempt to amplify the few real interactions that exist in a poor library. This observation again indicates that increases in random background signal may not always stem from actual random ligation in solution due to nuclei lysis, but can be caused by other factors.

## 4. Considerations in sequencing and mapping

### 4.1 How much can be gained by deeper sequencing?: Issues of library complexity

The potential number of pairwise genomic interactions in a mammalian genome is on the order of 1011 – 1012, so large numbers (hundreds of millions to a billion) of paired sequence reads are often needed to thoroughly sample this interaction space at high resolution. However, deeper sequencing will not lead to better Hi-C maps for all samples. The input number of cells and the efficiency of digestion and ligation limit the number of possible captured interactions. If HindIII digestion and ligation were perfectly efficient, 5 million cells could indeed lead to a library of 1012 captured interactions. However, inefficiencies and losses occur at many steps, so that this number in practice is often much more limited. Indeed, a highly complex replicate in which digestion, biotin fill-in, and ligation went well can provide better interaction maps with additional sequencing. However, a poorly complex replicate will barely change even when the number of reads is increased to 1 billion (Figure 4A). A high proportion (more than a few percent) of PCR duplicate molecules in a Hi-C library can also signal a lack of complexity. But, the increased prevalence of optical duplicates on patterned HiSeq 3000 and 4000 flowcells [42] adds uncertainty to this potential measure of complexity.

**Figure 4.**
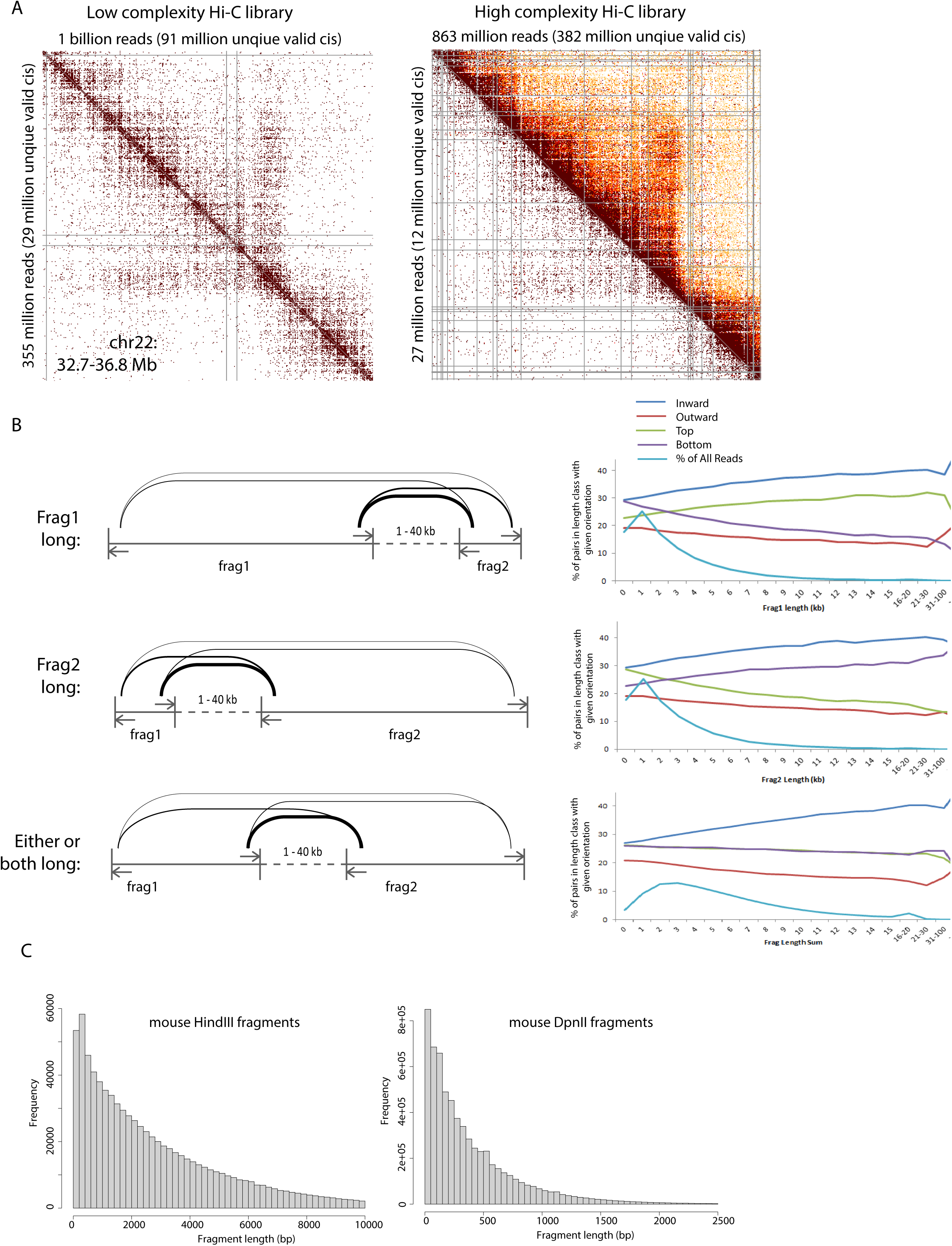
Considerations in Hi-C data alignment and binning. A) 10 kb binned and corrected heatmaps in a 4 Mb region of chr22 for varying input read numbers for poor complexity and good complexity replicates of a GM12878 Hi-C experiment. The poor complexity Hi-C library (left) gives no additional interaction resolution or detail even when input reads are increased to 1 billion (upper triangle) vs. 335 million (lower triangle). The high complexity library (right) on the other hand gives much more detailed information at 863 million reads (upper triangle), while only 27 million reads (lower triangle) are needed to give the amount of detail captured by 355 million reads of the poor complexity library. B) Three scenarios of proximal interactions are considered for their effect on interacting strand/side bias: “Frag1 long”: With increasing upstream restriction fragment length, positive strand interactions (“top”, green line) are increasingly favored compared to negative strand interactions (“bottom”, purple line). “Frag2 long”: with increasing downstream fragment length, the opposite pattern holds. “Either/both long”: if either fragment can be equally long, no strand bias is seen, only a preference for inward facing reads (closest on the linear genome) vs. outward facing reads (farthest apart on the linear genome). C) Distributions of HindIII (left) and DpnII (right) restriction fragment lengths in the mouse genome (mm9).

### 4.2 How finely can the interactions be binned?

#### 4.2.1 Structural information can be found in where along the fragment a read occurs

Interactions detected by sequencing Hi-C products are often mapped to the midpoint of each represented restriction enzyme, based on the idea that the crosslinked interaction could have occurred anywhere across the given fragment. We have found, however, that sometimes one end of the fragment is favored for an interaction over the other end, and that this bias can contain biologically significant information about the position of the specific interaction. For example, for interactions of between long fragments that are within 40 kb of each other along the linear genome, more ligations are observed between the fragment ends that are closer together (Figure 4B). This shows that the natural decay of interaction frequency along the chromatin polymer is revealed in a restriction fragment end preference for sequence reads. We therefore argue that it is valid, if convenient for other analyses, to bin sequence reads to constant bin sizes, taking into account exactly where each read lands rather than only which restriction fragment it belongs to. Employing this approach, however, absolutely requires that resulting interaction maps be carefully corrected for artifacts by a fragment-position naïve method such as iterative correction [34].

#### 4.2.2 Comparing restriction fragment size distributions to a chosen bin size

When choosing a high-resolution bin size for Hi-C data generated with a given restriction enzyme, it is also important to consider the restriction fragment size distribution to limit the number of bins containing no fragment ends. Publications often quote the average or median restriction fragment size, but we note that this can be a misleading value. HindIII and DpnII fragments follow a non-normal distribution (Figure 4C) in the human genome, so the average or median size does not imply that most fragments are close to this size. In fact, while the average HindIII fragment size in the mouse genome is 3.2 kb, nearly 30% of fragments are less than 1 kb in size, so there are numerous interactions that can be detected at a higher resolution than the average fragment size. Conversely, the median DpnII fragment size in the mouse genome is 260 bp, but nearly 30% of DpnII fragments are larger than 500 bp, so using a bin size of 200-300 bp to match the median fragment size will result in numerous bins with no information.

## 5. Considerations for applying Hi-C to lower input cell numbers

Interpreting Hi-C data always requires the consideration that the data represents the average structure across a population of cells [25]. Single cell Hi-C approaches have made it possible to capture interactions that occur in individual cells [43], but these datasets are inherently limited in their information, since a maximum of 4 interactions (2 ends of each fragment on 2 alleles) can be captured from each genomic location in any given cell. Therefore, there is a need for continued development of protocols effective for small populations of cells, particularly as Hi-C is applied to more primary tissues or to capture genome structure during rare cellular events. We have shown earlier that results between replicate batches of 5 million cells are highly reproducible (Figure 2), but many applications will require lower cell numbers than this. We have noted several experimental considerations that will be important as Hi-C is adapted to lower numbers of cells.

### 5.1 Steps prone to substantial DNA loss

With small numbers of cells, recovery of all possible DNA is important at every step. We have found unexpected losses of DNA at certain steps of the Hi-C protocol, which would be unacceptable if working with low cell numbers.

#### 5.1.1 Ethanol precipitation

Ethanol precipitation performed in large volumes as recommended by many Hi-C protocols can be inefficient. Indeed, we have evidence that substantial amounts of Hi-C DNA can be left behind in the ethanol precipitation of DNA during purification after the reversal of DNA crosslinks. In two independent experiments, we have performed another round of DNA precipitation on the ethanol supernatant removed from the first DNA purification attempt, and have recovered microgram quantities of additional DNA (10-40% of the original amount recovered) (Figure 5A). This indicates that in the first round of ethanol precipitation, substantial quantities of DNA were lost. Indeed, as 4C experiments are adapted to lower cell numbers, smaller volume DNA precipitations and minimizing tube transfers have been found to be valuable [44].

**Figure 5.**
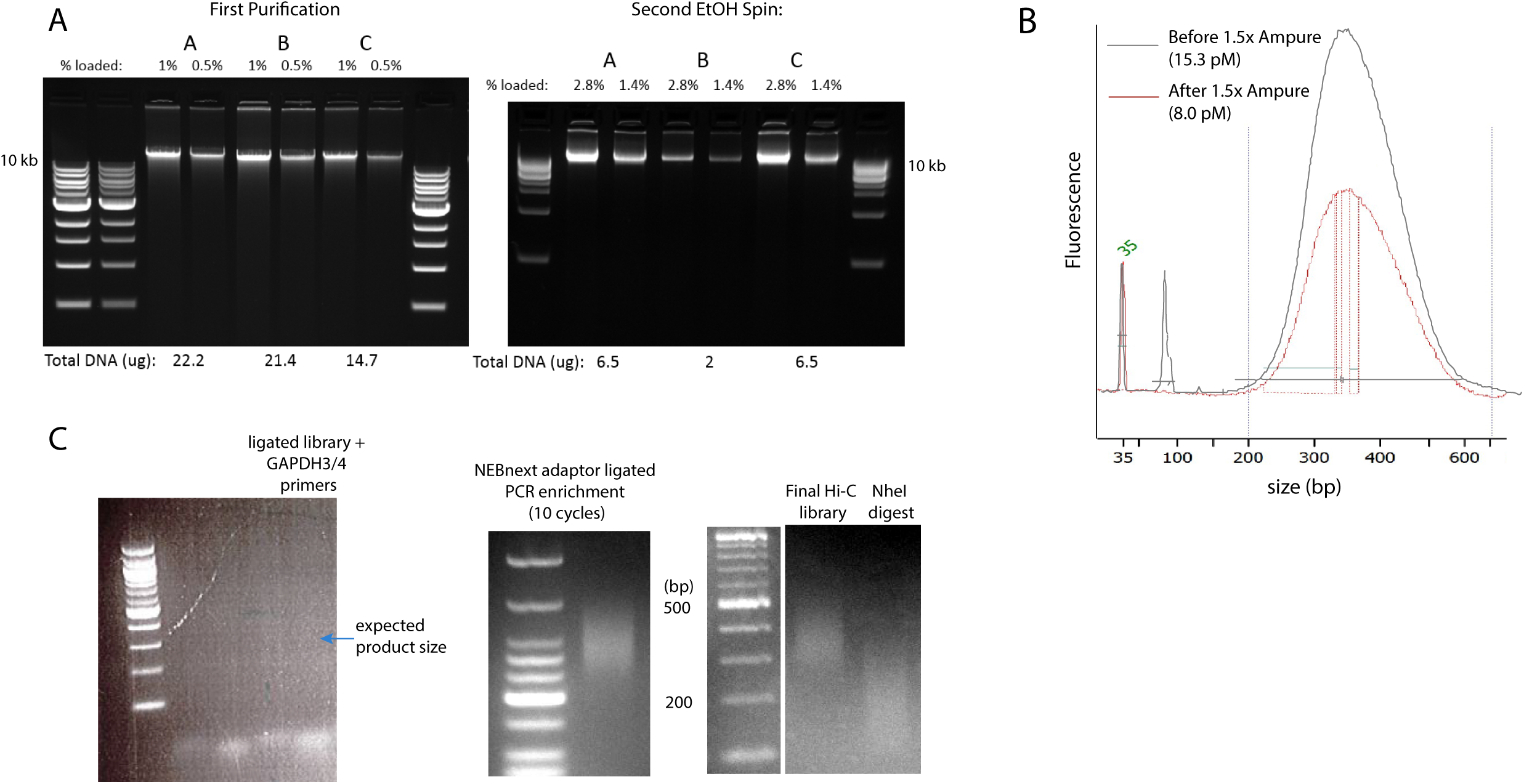
A) After reversal of crosslinks, three different GM12878 Hi-C libraries (A, B, and C) were purified and ethanol precipitated as recommended [31], yielding 14-22 micrograms of DNA. The ethanol supernatant removed from the DNA pellet was then re-centrifuged at the same speed as in the first purification, and another 2 – 6 micrograms (10-40% more) of DNA was recovered. B) Bioanalyzer traces before (black) and after (red) 1.5x Ampure XP purification are shown, scaled according to their 35 bp standard peak. Size-selection of Hi-C library DNA with 1.5x Ampure XP beads successfully removes the ∼80 bp excess adaptor peak (black), but reduces the overall concentration of DNA by nearly half. C) For the HindIII Hi-C 2.0 experiment on 5 million B16-F1 cells, no PCR product was observed for a neighboring interaction primer pair in the GAPDH gene region (primers described in [31]) (left). However, the final Hi-C library could be amplified with 10 cycles of PCR (middle), and digested well with NheI, as expected for a high quality Hi-C library (right). This final library produced good Hi-C interaction data (see Figure 2) despite the failed quality control neighboring interaction PCR. Note: some lanes in gels have been excised for clarity, but all lanes shown in contiguous images were run on the same gel.

#### 5.1.2 Reversing crosslinks

Many Hi-C protocols recommend reversing formaldehyde crosslinks at 65 degrees C overnight. We have observed that this lengthy high temperature step can easily result in DNA degradation if any traces of DNase enzymes are present. Extreme caution must be taken to avoid any DNase contamination before this step, and minimizing the length of this incubation may help reduce the likelihood of DNA degradation.

#### 5.1.3 Ampure purification

Ampure XP beads are a powerful and convenient way to select sequencing libraries for a given size range and to remove adaptor and primer dimers. However, if not performed carefully, this step can also result in substantial DNA loss in the desired size range as well. We have observed a 50% loss of Hi-C library material when a second round of Ampure purification was performed to remove excess adaptor dimers (Figure 5B). Troubleshooting guidelines provided with Ampure beads should be considered to avoid loss of material. Further, since any purification step will involve some DNA loss, protocols should be optimized to minimize the number of purification steps required.

### 5.2 Difficulties with assessing quality for Hi-C with low cell numbers

Previously published Hi-C protocols often recommend checking the quality of a Hi-C library after ligation and DNA purification by performing a targeted PCR on one or a few neighboring interaction products [24, 31]. However, we find that this control is less likely to be a reliable indicator of Hi-C library success when smaller input cell numbers are used. Though interaction patterns are reproducible as cell numbers are decreased from 20 million to 5 million (Figure 2B), the likelihood of detecting with PCR any given single interaction out of a small sample of the Hi-C library decreases with lower cell numbers. We have observed cases in which the quality control PCR of one interaction gave no product, but the final Hi-C library and sequenced results looked high quality (Figure 5C).

Indeed, even with larger numbers of cells, a PCR digest control can be unreliable. In a recent publication of Hi-C in barley, for example, two variations of Hi-C were used. Effective NheI digestion of the single interaction PCR product band suggested that one protocol worked much better than the other, but NheI digestion of the complex final library reveals that both experiments are similarly high quality [20]. In general, different quality control metrics may be needed to evaluate Hi-C experiments with low cell numbers. Experiments adapting 4C to low cell numbers have found, for example, that short range interaction profiles from low cell numbers can be reproducible even while weak long-range interactions are harder to detect reproducibly [44].

## 6. Conclusions

The complex nature of Hi-C experiments means that there are many steps, from cell harvesting to sequence mapping, which can affect the quality of the downstream genome interaction maps. Protocol improvements are continuing to increase the power, information, and resolution that can be gleaned from Hi-C experiments. Adjustments to the protocol such as those suggested in Hi-C 2.0, and ideas suggested here, such as avoiding over-sonication, can notably increase the recovery of informative interaction pairs. Artifacts and noise in the Hi-C data can arise from variables in the early cell preparation steps or as a result of the complex mixtures subjected to end repair and PCR steps. Some of these sources of artifacts still lead to occasional experiments with high levels of noise. Fortunately, as we have shown here, many features of Hi-C maps are reproducible despite protocol variations. As more scientists apply Hi-C to study 3D genome structures in diverse systems, careful attention to and documentation of the details of successful and problematic Hi-C experiments will lead to continued insights into how to optimize this important approach.

## Acknowledgements

We thank Yang Xu for his help in calculating Hi-C dataset reproducibility. We thank Job Dekker, Johan Gibcus, Jon Belton, Houda Belaghzal, Natalia Naumova, Noam Kaplan, and Bryan Lajoie for helpful discussions about concepts discussed here. Comparative Hi-C protocol experiments were in part supported by a Ralph E. Powe Junior Faculty Enhancement Award from Oak Ridge Associated Universities.

### Appendix A

#### Detailed protocols for in situ Hi-C and Hi-C 2.0 variations tested with HindIII digestion. NEBNext library preparation adaptations detailed (page 10) as well

This protocol is modeled on the Hi-C 2.0 protocol published by Belaghzal and Gibcus, 2017. The complete Hi-C protocol we used is included, with variations we tested for in situ Hi-C and Hi-C 2.0 noted. The 5 million cell aliquot used for the DpnII Hi-C 2.0 was processed according to the protocol below through step 2.3 and then exactly as described in Belaghzal and Gibcus, 2017.

### Day 1. CROSSLINKING OF CELLS, CELL LYSIS, AND CHROMATIN DIGESTION

*Grow the cells in appropriate culture medium. 5 million cells per Hi-C library are used. The degree of cell confluence, morphology, etc. should be documented carefully before each Hi-C experiment for future reference.*

### 1. CROSSLINKING ADHERENT CELLS (∼90 min)

*Volumes below are for a single T-75 Plate*

*Prepare 10 mL 1x HBSS + protease inhibitor cocktail per plate, place on ice*

1.1) Aspirate the medium, wash with 10 ml 1x HBSS (no serum) per plate.

1.2) Immediately before adding to the plate, mix 275 μl of 37% formaldehyde with 10 mL 1x HBSS to obtain 1% final formaldehyde concentration. Aspirate 1x HBSS wash from cell plate and add this pre-mixed 1% formaldehyde solution.

1.3) Incubate at room temperature (RT) for exactly 10 min on a gently shaking platform.

1.4) To quench the crosslinking reaction, add 554 μL of 2.5 M glycine, mix well

1.5) Incubate for 5 min at RT (rocking platform) and then incubate on ice for at least 15 min to stop crosslinking completely.

1.6) Aspirate liquid off of plates (discard as hazardous waste) and add 10 mL ice cold 1x HBSS + protease inhibitors

1.7) Scrape the cells from the plates with a cell scraper and transfer to a 50 ml tube.

1.8) Centrifuge aliquots of 5 million cells each at 800x*g* for 10 min, 4°C. Check to make sure all cells are pelleted. If there are visible floating cells, spin again.

1.9) Discard the supernatant by aspiration.

1.10) Cells can be snap-frozen in liquid nitrogen and stored at -80°C for at least 1.5 years or one can continue with cell lysis.

### 2. CELL LYSIS AND CHROMATIN DIGESTION (WITH HINDIII) (∼75 min)

2.1) Resuspend one crosslinked cell aliquot (∼5×106cells) in 1 ml of ice-cold lysis buffer (see Belaghzal et al., 2017 or Belton et al., 2012 for buffer formulations) containing 10 μl protease inhibitor cocktail (100x, Thermo)

2.1.1) Incubate on ice for 15 min.

2.1) Dounce homogenize the cells on ice with pestle A.

2.2.1) Slowly move pestle up and down 30 times

2.2.2) Incubate on ice 1 min to let the cells cool down

2.2.3) Then do 30 more strokes.

2.2.4) Image lysed cells under the microscope to check for intact nuclei and document cell appearance

2.3) Transfer the lysate to a 1.7ml tube

2.4) Centrifuge for 5 minutes at (in situ: 2,000xg /Hi-C2.0: 2,500x*g*) at RT to pellet nuclei.

2.5) Discard the supernatant and then wash the pellet twice by resuspending it in 500 μl of ice cold 1x CutSmart Buffer and then centrifuging the sample for 5 min at (in situ: 2,000xg /Hi-C2.0: 2,500x*g*).

2.6) Resuspend the pellet in 1x CutSmart Buffer, so that the total volume of the suspension is 360 uL. (Add 100 uL of buffer first, check total volume, and then add remaining amount necessary to get 360 uL total)

*Save 18 µl of lysate as a chromatin integrity control:*

*Add 50 μl of 1x CutSmart Buffer and 10 μl of Proteinase K (10 mg/ml). Incubate for 30 minutes at 65°C. Purify DNA by single phenol-chloroform extraction without ethanol precipitation. Add 1μl of RNAseA (1 mg/ml) to the aqueous phase and incubate for 15 min at 37°C. Store at -20 °C overnight and run on a 0.75% agarose gel at the same time as the digested DNA sample (see Day 2 below). The sample is good if DNA is either stuck in the well or runs as a single high molecular weight band (>23 kb)*

*After taking* 18µl *lysate control we have 342 µl* remaining *total volume.*

2.7) Add 38 μl of 1% SDS to the Hi-C tube (*380 μl total).* Mix carefully by pipetting up and down. Avoid making bubbles (0.1% SDS final).

2.8) Incubate at 65°C for 10 minutes exactly to open chromatin

2.9) Place tubes on ice immediately after

2.10) Add 43 μl of 10% Triton X-100 to the Hi-C-tube (*423 μl total*) to quench the SDS (1% Triton final). Mix gently by pipetting up and down Avoid making bubbles.

2.11) Add 12 μl of 10x CutSmart Buffer to the Hi-C tube to compensate for added components (*435 μl total before restriction enzyme*)

2.12) Add 400U (4 μl of 100,000 units/ml) HF HindIII to the Hi-C tube (*439 μl total*) Mix gently.

2.13) Digest the chromatin overnight at 37°C on a Nutator.

### Day 2. BIOTINYLATION, LIGATION, and CROSSLINK REVERSAL

#### DNA INTEGRITY / DIGESTION CHECK

Incubate tube at 65°C for 20 mins in order to deactivate the endonuclease enzyme. *After this incubation put the tubes on ice until cooled to room temperature This inactivation depends on the enzyme used. Please check enzyme datasheet*

Save 10 µl of lysate as a digestion control *(429 uL left; replace with 10 uL 1x CutSmart Buffer)*

Add 50 μl of 1x CutSmart Buffer and 10 μl of Proteinase K (10 mg/ml). Incubate for 30 minutes at 65°C. Purify DNA by single phenol-chloroform extraction without ethanol precipitation. Add 1μl of RNAseA (1 mg/ml) to the aqueous phase and incubate for 15 min at 37°C.

Check undigested and digested aliquots by running on a 0.75% gel, 75-100 V for 1 h. *(while gel is running, proceed with preparing biotinylation mastermix with everything but expensive Klenow and biotin reagents. If gel results look good, immediately proceed with biotinylation)*

### 3. BIOTIN FILL-IN

3.3) Prepare a Biotin Fill-in MasterMix as follows:

| FILL-IN MIX (ADD IN THIS ORDER) | PER TUBE |
| --- | --- |
| milliQ water | 2 uL |
| 10x CutSmart Buffer | 6 µl |
| 10 mM dATP | 1.5 µl |
| 10 mM dGTP | 1.5 µl |
| 10 mM dTTP | 1.5 µl |
| 0.4 mM biotin-14-dCTP | 37.5 µl |
| 5U/ µl Klenow | 10.0 µl |
| TOTAL | 60 µL |

3.4) Add 60 μl of the Fill-in master mix to the digested Hi-C chromatin *(should be 500 μl total; if some volume decrease occurs during digestion step above, bring total volume up to 500 uL with 1x CutSmart Buffer).*

3.5) Mix gently by pipetting up and down without producing any bubbles

3.6) Incubate the tubes at (in situ: 37°C for 2 hours / Hi-C2.0: 23°C for 4 hours) in a ThermoMixer (900 RPM mixing; 10 sec every 5 min)

3.6.1) Prepare ligation mixture (below) during incubation.

3.6.2) Place on ice immediately after incubation

### 4. BLUNT END DNA LIGATION

4.1.1) Prepare the 665 μl ligation mix during biotinylation:

| LIGATION MIX | PER TUBE |
| --- | --- |
| in situ: 10x ligation buffer [NEB] | 120 µl |
| Hi-C2.0: 5x ligation buffer [Invitrogen] | 240 µl |
| 10% Triton X-100 | 120 µl |
| 10 mg/ml BSA | 12 µl |
| T4 DNA ligase [Invitrogen] | in situ: 10 µl (5 U/µL)<br>Hi-C2.0: 50 µl (1 U/µL) |
| Water | To 700 µl |
| TOTAL | 700 µL |

*Note: we have tried the Hi-C 2.0 protocol with both a total of 50 U and a total of 250 U of ligase and find that additional ligase appears neither to help nor hurt the outcome.*

4.2) Add 700 μl of the mix to the Hi-C tube (*1,200 μl total*).

4.3) Incubate all tubes at 16°C for 4 hours in ThermoMixer with interval shaking (900 RPM mixing, 10 sec every 30 min).

### 5. REVERSE CROSSLINKING

5.1) Add 25 μl of 20 mg/mL proteinase K to the ligated Hi-C tube and incubate at 65°C overnight in the ThermoMixer

5.2) Add another 25 μl of proteinase K to the Hi-C tube and incubating an additional 2 hours (*1,250 μl total Hi-C)*

### Day 3. DNA PURIFICATION

#### 6. DNA PURIFICATION

6.1) Remove tubes from 65 C and allow to cool down.

6.2) Add 2.4 ml saturated phenol:chloroform (1:1, pH=8.0) to 2 × 15 ml tubes (label Hi-C PC1 and Hi-C PC2) for Hi-C

*Use fume hood, appropriate gloves, and labcoat when handling phenol:chloroform*

6.3) Extract the DNA:

6.3.1) Transfer Hi-C DNA into Hi-C PC1 tube

6.3.2) Vortex for 30 seconds to obtain a homogenous, milky solution

6.3.3) Centrifuge at 1500xg for 5 minutes (room temperature)

6.3.4) Carefully transfer the aqueous phase to Hi-C PC2. Repeat vortex and spin.

6.4) Transfer the aqueous phase with Hi-C DNA (*∼1.2 ml*) to a clean 15 ml tube (adequate for high-speed centrifugation at 16,000 xg)

6.4.1) Add 1x TLE (pH=8.0) up to 2 ml.

6.4.2) Add 1/10 volume of 3M sodium acetate, pH=5.2 (*200μl*) and vortex briefly.

6.4.3) Add 2.5 volumes of ice-cold 100% ethanol (*5 ml*) and mix well by inverting the tubes several times.

6.5) Incubate the tubes at -80°C for 45 min – 1 h

6.6) Centrifuge tubes at 16,000x*g* for 30 min at 4°C.

6.7) Decant the supernatant with extra caution as the pellet detaches from the tube wall quite easily and air dry the DNA pellets very briefly (∼1 min).

6.8.1) The pellet might become invisible while drying out

6.8) Dissolve Hi-C pellet in 450μl of 1x TLE (pH=8.0) and transfer to 2 mL, 30kD Amicon columns.

6.9) Wash the column at least 3 times with the following steps:

6.10.1) Centrifuge at 14,000xg in tabletop centrifuge for ∼5 minutes (room temperature)

6.10.2) Remove flowthrough (to a temporary save tube). DNA stays in the volume left in the column.

6.10.3) Add 450 μl TLE

6.10) On the last spin, spin for 10 minutes instead of 5 minutes at 14,000xg

6.11) After the last wash, flip the column onto a new Amicon container tube and spin at 1,000xg for 2 min to recover the DNA from the column

6.12) Add 1 μl of RNAseA (1 mg/ml) to the sample and incubate for 30 mins at 37°C.

*DNA samples can now be stored at -20°C.*

### 7. QUALITY CONTROL OF HI-C LIBRARIES

7.1) Make a 1:10 dilution of the Hi-C library and run 2μl and 6 μl on a 0.8% agarose gel for quality control at 100V for 1 hr.

7.2) Quantify amount of DNA by densitometry. Use several dilutions of the NEB 1kb DNA ladder as a standard curve to estimate the DNA concentration more accurately.

7.3) Measure the Biotin Incorporation by a PCR digest:

7.3.1) Make 1:20 dilution of Hi-C DNA (2.5 uL Hi-C + 47.5 uL H2O)

7.3.2) Dilute primers to 10 uM in TLE

7.3.4) Perform a PCR to amplify one neighboring interaction as follows:

| INGREDIENT | 1 REACTION |
| --- | --- |
| Diluted Template DNA | 20 uL |
| MILLIQ | 33.9 uL |
| 10X PCR buffer | 7.5 µl |
| 25mM dNTP | 0.6 µl |
| 50 mM MgSO4 | 6.0 uL |
| Primer 1 (10 µM) | 3 µl |
| Primer 2 (10 µM) | 3 µl |
| Taq DNA polymerase | 1 µl |
| Total | 75 uL |

7.4) Combine 20 uL diluted Hi-C DNA or water (neg ctrl) with 55 uL mastermix in a PCR tube.

7.5) Run the following PCR program:

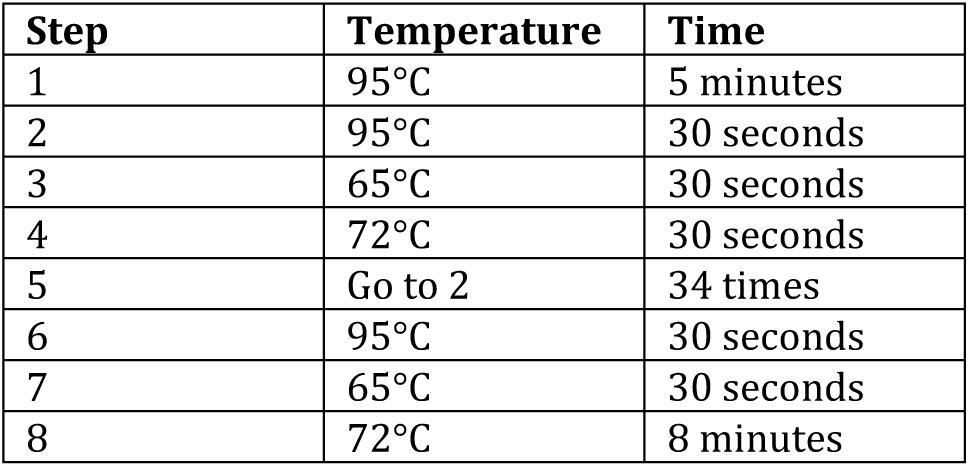

7.6) Run the samples in a 2% gel at 160 V for 45 min.

7.7) Digest the HiC PCR product with HindIII and NheI as follows:

7.6) Run the samples in a 2% gel at 160 V for 45 min. 7.7) Digest the HiC PCR product with HindIII and NheI as follows:
|  | Control (μl) | HindIII (μl) | NheI (μl) | HindIII/ NheI (μl) |
| --- | --- | --- | --- | --- |
| **PCR product** | 16.0 | 16.0 | 16.0 | 16.0 |
| **10X Cutsmart buffer** | 2.0 | 2.0 | 2.0 | 2.0 |
| **HindIII** | --- | 1 | --- | 1 |
| **NheI** | --- | --- | 1 | 1 |
| **MilliQ** | 2 | 1 | 1 | --- |

7.8) Incubate samples at 37°C for at least 30 minutes:

7.8.1) Combine each with 4 uL 6x dye and load 17 uL on the top row of wells in the 2% gel. Run 1 hr at 175 V.

Successful biotin fill-in of HindIII sites (AAGCTT) followed by blunt-end ligation create sites for the NheI restriction enzyme (GCTAGC).

### 8. REMOVAL OF BIOTIN FROM UN-LIGATED ENDS

8.1) Quantify DNA from Hi-C ligated library gel (step 7.2). Per 5 ug of DNA recovered, prepare reactions as follows:

| INGREDIENT | Per 5 ug |
| --- | --- |
| Hi-C DNA sample | **5 μg** |
| **10X NEBuffer 2.1** | 5 μl |
| 1 mM dATP | 1.25 μl |
| 1 mM dGTP | 1.25 μl |
| 3,000 U/ml T4 DNA polymerase | 5 μl |
| MILLIQ water | TO 50 μL |

8.1.1) Place each 50 μl reaction in a different well of a PCR strip

8.1.2) Incubate at 20°C for 4 hours in the PCR machine.

8.2) (in situ: Add 2 μl of 0.5M EDTA to each tube to stop the reaction ; Hi-C2.0: Inactivate the enzyme for 20 mins at 75°C)

8.3) Cool down/ keep at 4°C

8.4) Pool the reactions together and add water up to 500 uL. Perform an Amicon wash:

8.4.1) Add volume to Amicon column and centrifuge at 14,000xg for ∼ 5 min

8.4.2) Wash twice with 400 μl of milliQ water (spinning 5 min at 14k x g each time)

8.4.3) Invert the Amicon column onto a fresh tube and centrifuge for 2 minutes at 1,000xg to collect the DNA *Note that Amicon concentration does not remove proteins (they are only inactivated by the heat or EDTA treatment above, and then also by sonication below). Concentration by volume reduction may result in DNA coming out of solution. If this occurs, sonication below will help re-solubilize DNA.*

8.5) Add milliQ to a total of 132 μl. Save 2 uL for pre-sonicated gel check and transfer the rest to a Covaris microtube for sonication.

### 9. DNA SONICATION

9.1) Shear the DNA to a size of 200 – 400 bp using the Covaris M220 sonicator and the following parameters:

- 20% Duty Factor, 200 cycles/burst, 50 W Peak Incident Power for 110 seconds at 4 degrees C. (note that this differs from the Hi-C 2.0 recommended settings for reasons discussed in the manuscript)

9.2) Check the results of sonication on a 2% gel. Run the gel at 160 V for 45 min.

### 10. SIZE FRACTIONATION USING AMPURE XP

#### This step can be omitted for low concentrations of DNA and sonicated DNA can be taken directly to step 11

*The liquid phase of AMpure mixture precipitates DNA onto the AMpure XP magnetic beads. The size of the DNA molecules precipitated depends on the ratio of the volume of AMpure XP liquid (PEG8000) to the volume of the sample when mixed together. Increasing the proportion of AMpure XP liquid decreases the cutoff of the size of the DNA molecules precipitated onto the beads. Here, the 0.7x Ampure step removes fragments above ∼400 bp and the 1.2x Ampure captures the remaining fragments between 100-400 bp.*

Note that the 0.7x and 1.2x recommendation differs from Hi-C2.0 in order to capture the larger fragments coming out of sonication conditions above.

10.1) Bring volume of Hi-C sample up to 500ul with 1x TLE

10.2) Allow AMpure XP mixture to come to RT and vortex prior to use. Add 350 μl of AMpure XP mixture to the Hi-C tube. Label the tube as 0.7X *(the ratio of AMpure XP solution volume to Hi-C sample volume is 350/500 = 0.7).*

10.3) Vortex and spin down tubes briefly.

10.4) Incubate tubes for 10 min at RT.

10.5) Place tubes on the Magnetic Particle Separator (MPS) for 5 min at RT.

10.5.1) During above incubation, prepare a fresh tube with 500 μl AMpure XP mixture; label them as 1.2X.

10.5.2) Incubate tubes on the MPS for 5 min

10.5.3) Remove supernatant

10.5.4) Resuspend beads in 250 μl AMpure mixture

*This step increases the number of beads present in a smaller volume of AMpure XP mixture. This prevents saturation of the beads with DNA, ensuring an efficient precipitation of the DNA.*

10.6) Collect the 0.7X supernatants from MPS

10.6.1)Add the supernatants to the 1.2X tubes.

*The ratio of AMpure XP solution volume to the original sample volume is now: (250 + 350)/500 = 1.2. Under these conditions beads bind DNA fragments >100 bp.*

10.6.2) Vortex and spin down tubes briefly.

10.6.3) Incubate 1.2X tubes for 10 min at RT

10.6.4) Place 1.2X tubes on the MPS for 5 min at RT

10.7) Discard supernatant from the 1.2X tubes.

10.8) Wash the beads in 0.7X and 1.2X tubes twice with 1 ml fresh 70% ethanol, reclaiming beads against the MPS for 5 min each time.

10.9) Air dry beads on the MPS briefly (too much drying may decrease elution efficiency)

10.10) Resuspend the 0.7X and 1.2X beads in 150 μl of 1x TLE buffer in each tube to elute the DNA 10.10.1) Incubate 10 min at RT

10.10.2) Separate AMpure beads from eluate for both 0.7X and 1.2X tubes on the MPS for 5 min 10.10.3) Keep the eluate

10.10.4) For the gel, combine:

7 uL 6x dye + 10 uL 1.2x eluate + 10 uL water

2 uL 6x dye + 4.5 uL 0.7x eluate + 4.5 uL water

10.11) Desalt and concentrate the Hi-C sample on Amicon:

10.11.1) Centrifuge at 14,000xg for ∼5 min.

10.11.12) Add 450 μl of 1x TLE buffer

10.11.3) Centrifuge again until at 15,000xg for about 5∼10min until ∼30 μl remains

10.12) Place the Amicon unit upside down onto a new tube provided with the Amicon kit. Collect Hi-C sample by spinning at 1,000xg for 2 min

10.13) Bring volume of the sample to 50 μl with milliQ water

10.14) Check the quality and estimate the quantity of DNA on a 2% agarose gel. Run one lane of the 0.7X sample and a dilution series of the 1.2X sample along with a low molecular weight DNA marker. Quantify the amount of DNA in the 1.2X sample and compare to sonication gel to evaluate degree of DNA loss due to Ampure.

**NEBNext Protocol Adapted for Hi-C library Preparation on Beads**

At this step, the Hi-C 2.0 protocol recommends end repair and A tailing, then bead pulldown, followed by Illumina adapter ligation. We have adapted the entire NEBNext protocol to work after bead pulldown.

### 11. BIOTIN PULLDOWN WITH STREPTAVIDIN COATED BEADS

11.1) Vortex the MyOne™ Streptavidin C1 beads and then transfer 20 μl of beads to a 1.7ml LoBind tube.

11.2) Wash the beads with 400 μl of Tween wash buffer (TWB) by pipetting up and down and incubating for 3 min at RT on a rocking platform.

11.3) Reclaim beads against the MPS for 1 min, discard the supernatant.

11.4) Resuspend beads in 400 μl of TWB and transfer to a new LoBind tube. Incubate 3 min again. 11.5) Reclaim beads against the MPS for 1 min, discard the supernatant.

11.6) Resuspend beads in 400 μl of 2X Binding Buffer (BB) and add the DNA (in 400 μl TLE buffer)

11.7) Incubate the sample for 15 min at RT with rotation.

11.8) Reclaim the beads against the MPS for 1 min; discard the supernatant to a “BB sup” save tube.

11.9) Resuspend the beads in 400 μl of 1X BB and transfer them to a new tube

11.10) Reclaim the beads against the MPS for 1 min, discard the supernatant.

11.11) Wash beads with 100 μl of TLE and transfer to a new tube

11.12) Reclaim the beads against the MPS for 1 minute; discard the supernatant

11.13) Finally resuspend the beads in 50 uL TLE buffer.

### 12. NEBNext End Prep

12.1 Add the following components to a sterile nuclease-free tube:

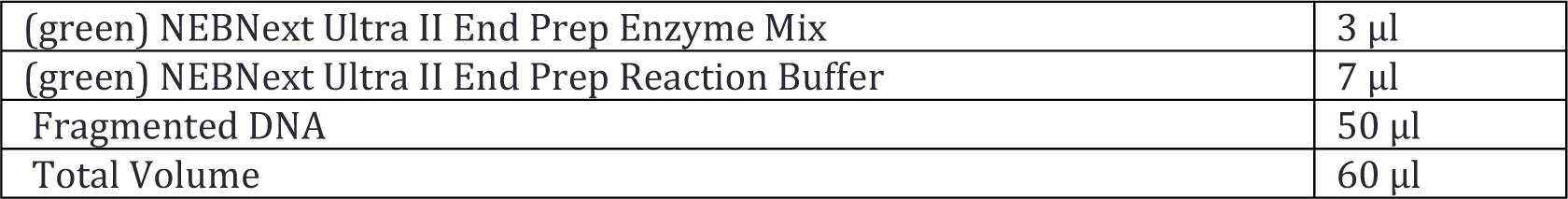

12.2 Set a 100 μl or 200 μl pipette to 50 μl and then gently pipette the entire volume up and down at least 10 times to mix throughly. Perform a quick spin to collect all liquid from the sides of the tube.

*Note: It is important to mix well. The presence of a small amount of bubbles will not interfere with performance.*

12.3 Place in a thermocycler, with the heated lid set to ≥ 75°C, and run the following program:

30 minutes @ 20°C

30 minutes @ 65°C

Hold at 4°C

*If necessary, samples can be stored at -20°C; however, a slight loss in yield (∼20%) may be observed. We recommend continuing with adaptor ligation before stopping.*

**13. Adaptor Ligation**

13.1 Perform a 1:15 or 1:20 dilution (or more if a first attempt gives excess adaptors) of the NEBNext Adaptor for Illumina in 10 mM Tris-HCI, pH 8.0 with 10mM NaCl.

13.2 Add the following components directly to the End Prep Reaction Mixture:

|  |  |
| --- | --- |
| End Prep Reaction Mixture (Step 12.3) | 60 µl |
| (red) NEBNext Ultra II Ligation Master Mix* | 30 µl |
| (red) NEBNext Ligation Enhancer | 1 µl |
| (red) NEBNext Adaptor for Illumina ** | 2.5 µl |
| Total volume | 93.5 µl |

\* Mix the Ultra II Ligation Master Mix by pipetting up and down several times prior to adding to the reaction.
*Note: The Ligation Master Mix and Ligation Enhancer can be mixed ahead of time and is stable for at least 8 hours @ 4°C. We do not recommend adding adaptor to a premix in the Adaptor Ligation Step.*

13.3 Set a 100 μl or 200 μl pipette to 80 μl and then pipette the entire volume up and down at least 10 times to mix thoroughly. Perform a quick spin to collect all liquid from the sides of the tube.

*(Caution: The NEBNext Ultra II Ligation Master Mix is very viscous. Care should be taken to ensure adequate mixing of the ligation reaction, as incomplete mixing will result in reduced ligation efficiency. The presence of a small amount of bubbles will not interfere with performance).*

13.4 Incubate at 20°C for 15 minutes in a thermocycler with the heated lid off.

13.5 Add 3 μl of (red) USER™ Enzyme to the ligation mixture from Step 13.4.

*Note: Steps 13.5 and 13.6 are only required for use with NEBNext Adaptors. USER enzyme can be found in the NEBNext Singleplex (NEB #E7350) or Multiplex (NEB #E7335, #E7500, #E7600, #E7710, #E7730 and #E6609) Oligos for Illumina.*

13.6 Mix well and incubate at 37°C for 15 minutes with the heated lid set to ≥ 47°C.

*Note: Samples can be stored overnight at -20°C.*

13.7) Reclaim the beads against the MPS for 1 min. Discard the supernatant.

13.8) Wash the beads twice with 400 μl of TWB. Add buffer to the beads, mix carefully by pipetting and incubate at RT for 5 min with rotation.

13.9) Reclaim the beads against the MPS for 1 min. Discard the supernatant. (to a save tube)

13.10) Resuspend the beads in 200 μl of 1X BB and transfer to a new tube

13.11) Reclaim the beads against the MPS for 1 minute. Discard the supernatant.

13.12) Wash the beads twice with 200 μl TLE. Resuspend beads then transfer to a new tube

13.13) Reclaim the beads against the MPS for 1 min. Discard the supernatant.

13.14) After the last wash, resuspend the beads in 20 μl of TLE and transfer to a new tube.

*Note: Samples can be stored at -20°C.*

**14. PCR Enrichment of Adaptor-ligated DNA**

*Note:Check and verify that the concentration of your oligos is 10 μM.*

14.1. PCR TITRATION and PRODUCTION PCR

14.1.1) Use the following HiCPCR_6CYC and HiCPCR_3CYC programs for PCRs below:

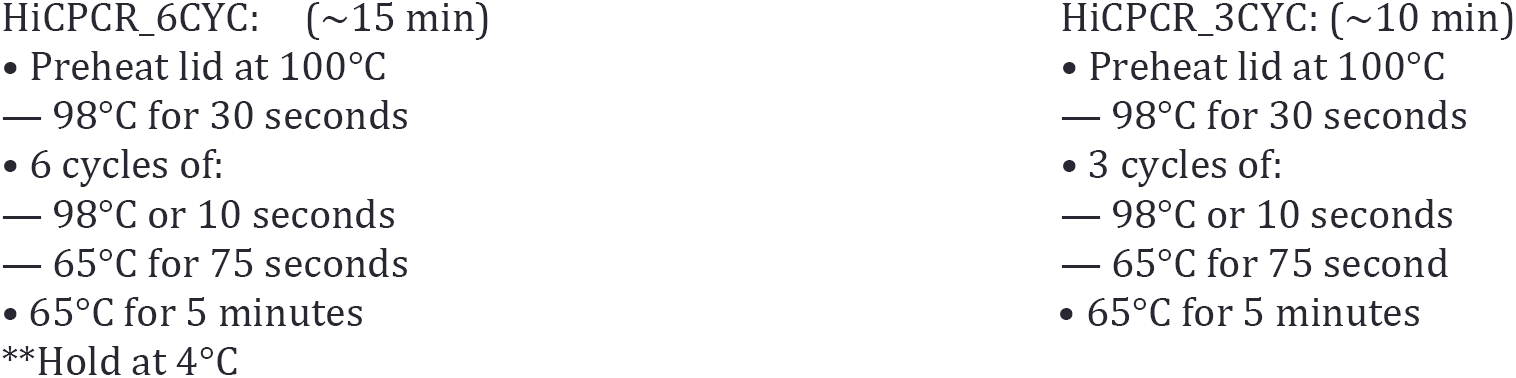

14.1.2. Add the following components to a sterile strip tube:

14.1.2A. **Forward and Reverse Primer <u>not</u> already combined**

|  |  |
| --- | --- |
| Adaptor Ligated DNA Fragments (Step 13.14 ) | 3 µl |
| (blue) NEBNext Ultra II Q5 Master Mix | 12.5 µl |
| (blue) Index Primer/i7 Primer*, ** | 2.5 µl |
| (blue) Universal PCR Primer/i5 Primer*, *** | 2.5 µl |
| Add water to Total Volume of: | 25 µl |

* The primers are provided in NEBNext Singleplex (NEB <u>#E7350</u>) or Multiplex (NEB <u>#E7335</u>, <u>#E7500</u>, <u>#E7710</u>, <u>#E7730</u>, <u>#E7600</u>) Oligos for Illumina. For use with Dual Index Primers (NEB <u>#E7600</u>), look at the NEB <u>#E7600</u> manual for valid barcode combinations and tips for setting up PCR reactions.

** For use with NEBNext Multiplex Oligos (NEB <u>#E7335</u> or <u>#E7500</u>, <u>#E7710</u>, <u>#E7730</u>) use only one index primer per PCR reaction. For use with Dual Index Primers (NEB #E7600) use only one i7 primer per reaction.

*** For use with Dual Index Primers (NEB <u>#E7600</u>) use only one i5 Primer per reaction.

14.1.2. Set a 100 μl or 200 μl pipette to 40 μl and then pipette the entire volume up and down at least 10 times to mix thoroughly. Perform a quick spin to collect all liquid from the sides of the tube.

Run the following PCR programs. (Note that this must be run when you have time to take aliquots every ∼10 min for 1 hour)

Set up 5 small tubes for gel loading labeled 6, 9, 12, 15, 18 and combine 2 uL 6x dye + 7 uL water in each.

After each PCR below, take out tube in the last 10 seconds of the 65 C final step, put tube directly on ice, and remove a 3 uL aliquot. Combine each aliquot with dye and water in tubes prepared above.

1. HiCPCR_6CYC (total 6 cycles)
2. HiCPCR_3CYC (total 9 cycles)
3. HiCPCR_3CYC (total 12 cycles)
4. HiCPCR_3CYC (total 15 cycles)
5. HiCPCR_3CYC (total 18 cycles)

14.1.4) Run the titration PCR on a 2% gel. Run the gel at 160 V for 45 min

14.1.5) Build a calibration curve showing the DNA quantity vs. number of cycles.

Choose an optimal number of cycles and number of PCR reactions for the final amplification of the library for deep sequencing. The cycle number should be chosen so that the PCR amplification is in the linear range and the expected size distribution is preserved (over cycling will shift the size distribution of the library towards higher molecular weight products). The number of PCR reactions should be calculated to produce the amount of DNA desired for sequencing (usually 50-100 ng). It is always better to reduce the number of PCR cycles while increasing the number of PCR reactions. The optimal amount is usually 1-2 cycles below the lowest amount visible on the gel.

### 15. PRODUCTION PCR

15.1) Decision from titration:_____ Perform PCR reactions (maximum of 5) with _____ cycles.

15.2) Set up N PCR master mix reactions on ice as follows:

|  |  |
| --- | --- |
| Adaptor Ligated DNA Fragments (Step 3.1.14 or 3.2.11) | 3 µl |
| (blue) NEBNext Ultra II Q5 Master Mix | 12.5 µl |
| (blue) Index Primer/i7 Primer*, ** | 2.5 µl |
| (blue) Universal PCR Primer/i5 Primer*, *** | 2.5 µl |
| Add water to Total Volume of: | 25 µl |

15.3) Run the following PCR program with the selected number of cycles.

- Preheat lid at 100°C
  – 98°C for 30 seconds
- N cycles of:

–98°C or 10 seconds
–65°C for 75 seconds
- 65°C for 5 minutes

15.4) Pool all PCR reactions together in a 1.7 mL Lo-Bind Tube and incubate on MPS for 2 min.

14.10.1) Remove supernatant to a new tube (this contains amplified DNA)

14.10.2) Resuspend beads in N*3 uL TLE and freeze at -20 in case of future need.

14.10.3) Take a 3 uL “pre-purified” aliquot of supernatant + 2 uL dye + 7 uL H2O for gel check.

15.6) To remove primer dimers, purify the amplified Hi-C library from the supernatant using AMpure XP beads.

15.6.1) Allow AMpure XP mixture to come to RT and vortex prior the use.

15.6.2) Add 1.5x volumes (uL) of AMpure XP mixture to the supernatant.

*Note: if the Bioanalyzer trace at the end of step 16 indicates high amounts of adapter-dimer or primer-dimer, re-purifying with 1.3x or 1.2x Ampure to remove those products is recommended.*

15.6.3) Vortex and spin down briefly.

15.6.4) Incubate for 10 min at RT.

15.6.5) Place on the MPS for 5 min at RT. Discard supernatant to a save tube.

15.6.6) Wash the beads twice with 1 ml of freshly made 70% ethanol

15.6.7) Air-dry the beads briefly and then resuspend in 35 μl of TLE buffer

15.6.8) Incubate the beads for 10 min at RT, tapping the tube every 1-2 min.

15.6.9) Collect the beads with the MPS for 5 min

15.6.10) Transfer the supernatant, containing the final Hi-C library, to a new tube.

Check final HiC library on a 2% agarose gel at 160 V for 45 min.

15.7) QUALITY CONTROL OF HI-C LIBRARY BY RESTRICTION DIGEST

16.1) Digest a small aliquot of the final Hi-C library with NheI to estimate the portion of molecules with valid biotinylated junctions.

|  |  |
| --- | --- |
| INGREDIENT | 1X |
| 10x NEB Cutsmart buffer | 1 µl |
| milliQ water | 6.5 uL |
| NheI-HF | 1 µl |
| Hi-C product | 1.5 uL |
| TOTAL | 10 μL |

16.2) Allow to digest at 37°C for at least 1 hour

16.3) Combine entire volume of digestion reaction + 2 uL dye and 1.5 uL uncut Hi-C DNA + 8.5 uL H2O + 2 uL 6x dye and run on a 2% agarose gel at 160 V for 45 min

16.4) Quantify approximate % of uncut DNA (unshifted smear) as a rough measure of potential fraction of non-ligation products in final sample

Run 1 uL of final Hi-C library on Agilent DNA Bioanalyzer. Dilute an aliquot of the library to 10 nM in 10 uL with Tris-HCl 10 mM, pH 8.5 + 0.1% Tween 20 for later pooling and sequencing.

